# Modelling novelty detection in the thalamocortical loop

**DOI:** 10.1101/2021.11.08.467674

**Authors:** Chao Han, Gwendolyn English, Hannes P. Saal, Giacomo Indiveri, Aditya Gilra, Wolfger von der Behrens, Eleni Vasilaki

**Author notes:** These authors contributed equally to this work (Wolfger von der Behrens, Eleni Vasilaki).

## Abstract

In complex natural environments, sensory systems are constantly exposed to a large stream of inputs. Novel or rare stimuli, which are often associated with behaviorally important events, are typically processed differently than the steady sensory background, which has less relevance. Neural signatures of such differential processing, commonly referred to as novelty detection, have been identified on the level of EEG recordings as mismatch negativity and the level of single neurons as stimulus-specific adaptation. Here, we propose a multi-scale recurrent network with synaptic depression to explain how novelty detection can arise in the whisker-related part of the somatosensory thalamocortical loop. The architecture and dynamics of the model presume that neurons in cortical layer 6 adapt, via synaptic depression, specifically to a frequently presented stimulus, resulting in reduced population activity in the corresponding cortical column when compared with the population activity evoked by a rare stimulus. This difference in population activity is then projected from the cortex to the thalamus and amplified through the interaction between neurons of the primary and reticular nuclei of the thalamus, resulting in spindle-like, rhythmic oscillations. These differentially activated thalamic oscillations are forwarded to cortical layer 4 as a late secondary response that is specific to rare stimuli that violate a particular stimulus pattern. Model results show a strong analogy between this late single neuron activity and EEG-based mismatch negativity in terms of their common sensitivity to presentation context and timescales of response latency, as observed experimentally. Our results indicate that adaptation in L6 can establish the thalamocortical dynamics that produce signatures of SSA and MMN and suggest a mechanistic model of novelty detection that could generalize to other sensory modalities.

**Author summary:** Cortical sensory neurons have been shown to be capable of novelty detection, that is they respond more vigorously when a novel, unexpected stimulus is presented, and less so when the stimulus is part of a predictable sequence. However, the neural mechanism underlying this capability is not yet fully understood. Here, we developed a thalamocortical network model that accounts for novelty detection and reproduces physiologically observed neural response patterns in the somatosensory cortex. Specifically, our results demonstrate that the novelty signal arises from the complex recurrent interplay between thalamic neurons and cortical neurons in layers 4 and 6. This work therefore provides a concrete mechanism that can serve as a starting point for further investigating the neural circuit mechanisms underlying novelty detection.

## Introduction

Sensory cortices are adept at identifying regularities and patterns obscured in the ever-changing input stream to dynamically generate predictions for the forthcoming stimulus. Their responses are then updated according to differences between the anticipated sensory inputs and the actual ones [1]. Mismatch negativity (MMN), observed in human electroencephalograph (EEG) recordings [2], and stimulus-specific adaptation (SSA), evident in single-cell recordings in animals [3], are two prominent instances of this type of short-term plasticity acting over timescales of seconds to minutes. For example, when consecutively presented with an identical stimulus over a short period, the cortical sensory neuron will decline/adapt its response and predict/expect another same one for the next stimulus. If the next stimulus violates the expectation, a larger/unadapted response will be evoked than by the expected stimulus. Therefore, both SSA and MMN conceptually represent a form of novelty detection: unexpected or infrequent stimuli are made more salient by eliciting stronger responses than expected or frequent ones.

The phenomenon of attenuated neuronal responses to common (standard) stimuli without generalizing to other, rare (deviant) stimuli has been a long-standing topic of interest, especially in the auditory system, which is referred to as SSA [4–8]. It is typically tested using the oddball paradigm that contains a repetitive standard sequence occasionally interrupted by a deviant [9]. SSA is a robust and widespread phenomenon found in multiple sensory modalities. Ulanovsky et al [4] first observed this phenomenon in the auditory cortex, and following this pioneering work, SSA has also been identified at subcortical stages of the auditory pathway such as the thalamus [10–12] and the inferior colliculus [13, 14], as well as in other sensory systems such as the somatosensory cortex [5, 8] and the visual system [15, 16]. Furthermore, SSA was demonstrated in the auditory cortex of anaesthetized [7, 17], awake [18] and freely moving rats [19], indicating that this phenomenon is not significantly affected by brain state and likely a hardwired component of sensory processing.

MMN, like SSA, is usually studied by applying the oddball paradigm and manifested in human EEG recordings as an additional negative deviation elicited by a deviant stimulus that breaks the regularity established by repeated presentations of the standard [9]. Both cortical SSA and MMN demonstrate a “true” deviance-detecting property, where the deviant response is dependent on its presentation context and beyond mere sensory input depression by applying several control paradigms [6, 8, 20]. In addition to SSA, animal models also show MMN-like responses that are homologous with human MMN in terms of their electrophysiological, pharmacological and functional properties [21, 22]. Despite significant similarities, SSA is unlikely to be the direct neuronal substrate of MMN, but may be one of the several mechanisms leading to the generation of MMN [23]. While MMN has been pharmacologically demonstrated to be NMDA-receptor dependent [24], SSA seems to be not affected by NMDA-receptor antagonists [25]. Besides, the response latency of SSA (peaks <10 ms after stimuli onset) is substantially shorter than MMN (peaks ~ 150-200 ms post-stimulus). Instead, SSA temporally matches better another event-related potential called mid-latency response (MLR) that peaks around 20 ms after stimuli onset [22]. MLR observed in humans and animals also demonstrates true deviance detection [26] and its insensitivity to NMDA-receptor antagonists [27]. Thus, it seems more plausible to postulate that SSA is the potential neuronal correlate of mid-latency response. Moreover, Musall et al. [8] recently found a late deviant-sensitive sensory response in the primary somatosensory cortex (S1) occurring hundreds of milliseconds post-stimulus, similar to MMN latency. The late sensory response has also been tested to be context-specific. These evidence strongly implied that the late response possibly lies in the physiological origin of MMN.

Over the past decade, a large number of computational models have been proposed to account for cortical SSA, with a focus on the auditory pathway. One simple type of SSA model is based on a feedforward network with synaptic depression [17, 28–31] which, unlike general neuronal fatigue, is stimulus-specific. Prominent synaptic depression is found in the thalamocortical pathway, and thus cortical SSA emerges due to the differentially adapted thalamocortical inputs [17, 28]. In this class of models, the standard’s regularity cannot affect the deviant response because the responses to each stimulus feature solely depend on the amount of adaptation load accumulated in its input channel. Hence the model fails to exhibit true deviance detection in which the presentation context formed by the standards also contributes to the deviant response. Nevertheless, this flaw of the model can be addressed through some modifications, such as employing the model in a multi-layer configuration [30] or assuming the width of the tuning curve of the input channel is history-dependent [3]. On top of that, recurrent networks have been considered an alternative way to generate cortical SSA [32–36]. Predictive coding based neuronal network of the auditory cortex was studied by Wacongne et al. to account for deviance detection in the context of MMN [32]. Yarden et al. proposed a generic SSA mechanism mediated via the propagation of synchronous population activity across a local neural circuit [35]. Recently a laminar network of morphologically plausible, multi-compartmental neuron units was designed to capture SSA in the form of local field potentials [36].

However, there is still no computational model accounting for the biphasic sensory responses in S1 that can be either sensitive to the rarity and presentation context of stimuli or not, depending on the response latency [8]. In particular, the latency of the secondary cortical response cannot be accounted for by the dynamics of a recurrent network involving short-term synaptic depression that is commonly regarded as a indispensable constituent of SSA models. Thus the latency is possibly produced by some subcortical source(s) other than the intra-cortical dynamics. Inspired by the idea that SSA arises from the interaction between differentially adapted populations of neurons tuned for specific feature stimuli [35], here we developed a hierarchal thalamocortical model for the whisker-related region of S1 (also known as the barrel cortex) with multiple timescales of synaptic and neuronal dynamics to capture the biphasic activity that demonstrates both signatures of SSA and MMN. On the short timescale of the early response, a vital element of the transient dynamics of the cortical circuitry is the so-called population spike (PS), characterized by nearly coincident firings of a pool of neurons evoked by a brief stimulus. On the longer timescale, we hypothesize that the late rhythmic cortical response is inherited from the thalamic spindle oscillation initiated by the interplay between the thalamic reticular (RE) and thalamocortical (TC) relay cells. The somatosensory thalamocortical network retains only those crucial neurobiological features, such as the laminar architecture and somatotopic organization of the barrel cortex, synaptic connectivity between the barrel cortex and its corresponding thalamic nuclei, as well as different synaptic receptors within thalamic nuclei, that are sufficient to elicit the late context-dependent deviant responses. Finally, we test the cortical capacity for predictive coding using more complex stimulus sequences. The results of our model indicate that the precision of expectation generated by sequence history impacts the neural response to stimuli that interrupt those expectations.

## Results

The architecture of the somatosensory thalamocortical network is organized in a loop as shown in Fig 1A. Based on the stereotypic somatotopic map for representing rodent whiskers, the barrel cortex is modeled as a grid of interconnected vertical columns, each of which primarily mediates stimuli from its principal whisker. Also, we only focus on neuronal clusters within layers 4 (barrels) and 6 (infrabarrels) of each cortical column, which respectively act as the afferent and efferent layers of the cortex from and to the thalamus [37–39]. The barreloids in the ventral posterior medial (VPM) nucleus of the thalamus show an identical layout to that of the barrels. Each barreloid relays the information from its principal whisker through an ascending sensory pathway to L4 of its principal as well as surrounding barrels, with stronger coupling to the principal than the surrounding ones. We hypothesize that L2/3 and L5 contribute little to the cortical SSA but merely transmit columnar information from L4 to L6. Therefore, a direct shortcut connection from L4 to L6 within every cortical column is added, which is sufficient in the simplified model proposed here to demonstrate SSA in L6, although in reality the flow of excitation from L4 will go through L3, L2 and L5 in sequence before arriving at L6 [40–42]. To close the thalamocortical loop, the outputs of layer 6 are projected back to their somatotopically aligned barreloids and thalamic reticular population. Here we assume that neurons in the thalamic reticular nucleus (TRN) are also whisker-specific, but they are not necessarily arranged in spatially discrete clusters as barreloids.

**Fig 1.**
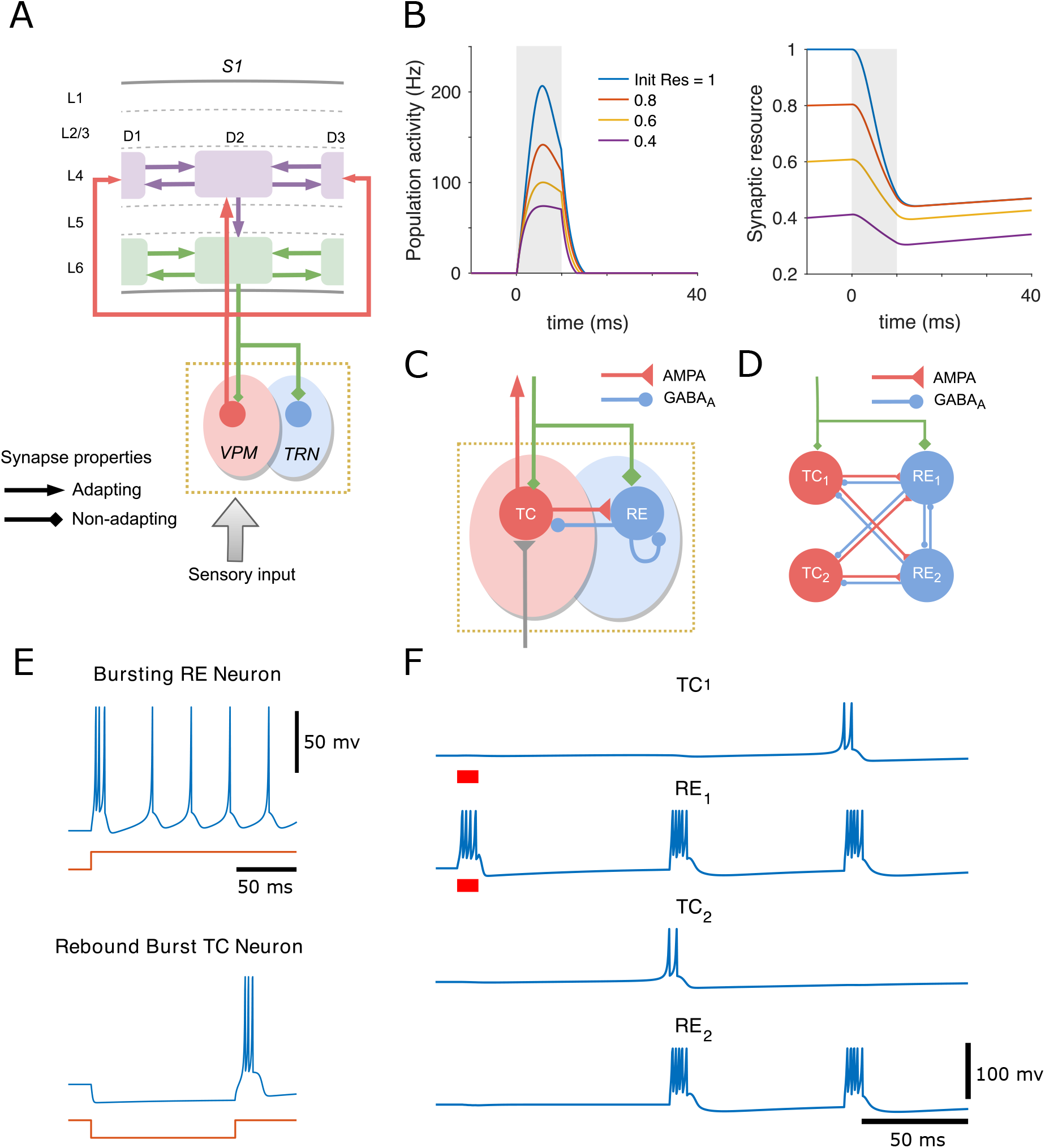
Structure and dynamics of the thalamocortical network and minimal thalamic circuit inducing spindle oscillations. (A) The ventral posterior medial (VPM) nucleus of the thalamus (pink shade), which is somatotopically arranged into discrete clusters called “barreloids”, relays peripheral sensory information to the L4 barrels (purple), with the relative strength of connections represented by the size of the arrows. Each L4 cluster projects its output vertically to the corresponding L6 infrabarrel (green), which in turn provides feedback excitation to its somatotopic barreloid and thalamic reticular nucleus (TRN, in blue shade), with higher cortical drive on TRN than VPM (specified by the size of the diamonds). Each L4 barrel and L6 infrabarrel also connects to its neighbouring barrels. (B) Population spike (PS) can be generated in the cortical cluster, represented by the transient increase of the cluster’s mean firing rate (left). Mean synaptic resources of the neuronal group are depleted by the PS and later recover gradually (right). Higher levels of initial resources evoke a more substantial PS. The time 0 represents the onset of a stimulus whose duration is marked in grey shading. (C) Two classes of cells with different synaptic receptors are used in the thalamic circuit: thalamocortical (TC) relay cells with excitatory AMPA-mediated synapses in the VPM and reticular (RE) cells with inhibitory GABA_A_-mediated synapses in the TRN. TC and RE neuronal populations are mutually coupled, and the RE population is also recurrently connected. The thalamus can be activated by bottom-up sensory input (grey) and top-down cortical drive (green). (D) A minimal thalamic network of 2 TC and 2 RE cells driven by the cortex is capable of inducing spindle oscillations. (E) Voltage traces of two types of thalamic cells simulated by the izhikevich model in response to a step in direct current (bottom of each panel): initial burst tonic firing in TC neuron (−70 mV at rest, hyperpolarizing pulse of 10 pA) and rebound burst in RE neuron (−62.5mV at rest, depolarizing pulse of −10 pA lasting 120 ms). (F) The time course of membrane potentials for the 4 thalamic cells. Cortical stimulation is marked by red bars under the traces (0.08 pA for TC_1_ and 20 pA for RE_1_).

We first focus only on the cortical network that is capable of generating SSA and true deviance detection by modulation of cross-barrel synchronous activities. All cortical intra- and inter-column synapses exhibit short-term depression. Only excitatory populations are considered in the cortical circuitry to simplify the mean-field analysis, and the level of excitability can be regulated by the threshold of neuronal gain function. Every layer-specific population in columns is modelled by a mean-field recurrent network coupled with depressing synapses, which can generate the so-called population spike (PS) in response to external stimulation. The PS is characterized by nearly synchronous firing of a group of neurons within a short time window [43, 44]. In the cortical mean-field rate populations we studied here, a population spike is described as a sharp increase in population activity, leading to fast depletion of mean synaptic resource that will recover gradually after the removal of external input. Higher initial synaptic resources cause more substantial PSs to be triggered in response to the same stimulus (Fig 1B).

The thalamic circuitry is set up to induce spindle oscillations characterized by intermittent bursting activity at 7-14 Hz, which is postulated as the origin of the late rhythmic responses in L4 of S1 [8], since the frequency of cortical oscillation (~10Hz) falls into the spindle range. Early studies have shown that network and intrinsic mechanisms act in combination to generate spindle oscillations [45]. Two interacting populations of spiking thalamic neurons are considered to model spindle oscillation: excitatory thalamocortical (TC) neurons and inhibitory thalamic reticular (RE) neurons. As illustrated in Fig 1C, the TC population is excited by whisker stimulation and then provides AMPA-mediated excitatory synaptic input to the RE population and neurons in cortical L4. In turn, the RE neurons suppress the activity of the TC neurons and themselves via GABA_A_-mediated inhibition. The top-down modulation from L6 exerts more effective excitation on RE than TC population due to more substantial corticothalamic conductance [38, 45, 46]. Unlike the mean-field dynamics of the cortex represented by the rate network, the intrinsic dynamics of individual spiking thalamic neuron is simulated by the izhikevich model (see Materials and methods section for model details). RE neurons exhibit transient bursting followed by tonic firing in response to depolarizing step currents, and TC relay neurons can trigger rebound spike-bursts upon release from hyperpolarization [45, 47] (Fig 1E).

We elucidate the mechanism of spindle oscillation triggered by the corticothalamic feedback excitation in a minimal thalamic circuit consisting of 2 RE and 2 TC cells [46] (Fig 1D). When the cortical drive is exerted on both TC_1_ and RE_1_ neurons, only the RE1 neuron is able to elicit spikes due to the considerably higher cortical conductance on RE than TC neurons. The bursting activity of RE_1_ cell provide inhibitory postsynaptic potentials (IPSPs) to both TC cells and then induce rebound bursting spikes in TC_2_ cells when its potential is released from hyperpolarization and almost reaches its resting state. The firing of TC_2_ cell reciprocally excite RE cells, leading to the next cycle of spindle oscillations (Fig 1F). Minute bump in the potential of TC_1_ cell evoked by initial corticothalamic stimulation prevents the cell from generating rebound bursts in the first cycle, which results in the TC cells fire every two cycles while RE cells fire every cycle in the minimal circuit. With the thalamocortical pathway, spindle oscillation is synchronized over cortical areas as hypothesized to be the late oscillatory rhythms found in L4 of S1 [8].

### Propagation of population activities in the thalamocortical loop

The PS evoked in the principal L4 barrel will propagate to the neighbouring L4 barrels via inter-column coupling, as well as to the column-aligned L6 infrabarrel via the L4-to-L6 projection [40]. The L4 responses are largely localized around the principal barrel and substantially attenuated in the surrounding barrels [48, 49] (Fig 2A, top panel). Responses across L4 barrels start almost simultaneously about 7 milliseconds after stimulus onset, in accordance with experimental findings [50] (Fig 2B, top). The topographical spread of PS in L6 is similar to that in L4 but with broader temporal profiles of the PSs and slightly longer response latencies [50] (Fig 2A and B, bottom panels). The extent of propagation is determined by the width of the cross-whisker tuning curve of thalamocortical input, the strength of inter-column connections as well as the excitability of the neural population, where a broader tuning curve, stronger inter-population couplings and a more excitable population give rise to more extensive propagation.

**Fig 2.**
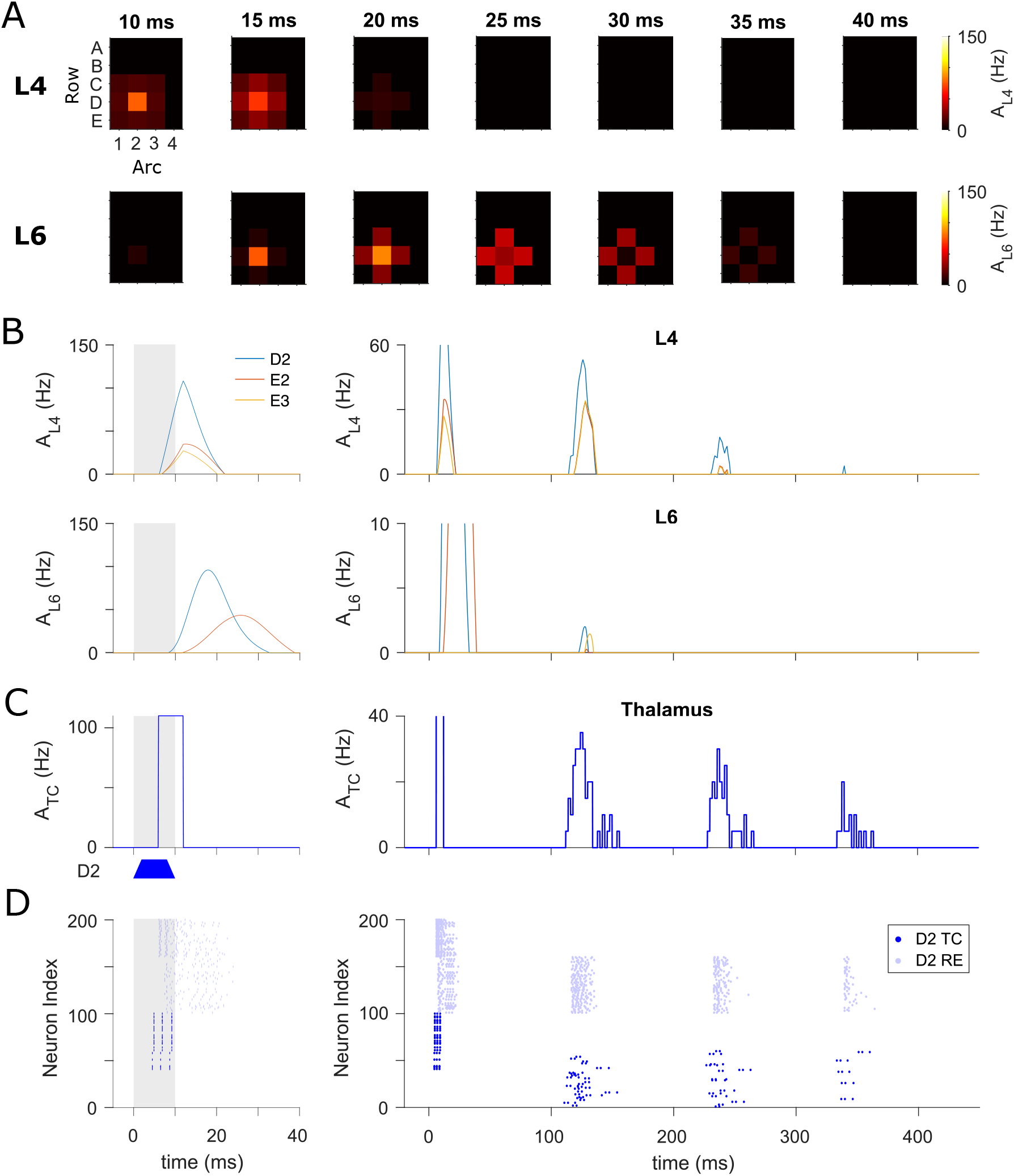
Spatiotemporal distribution of whisker-evoked responses in cortical columns and thalamus. (A) Top: Snapshots of cortical spatial activity patterns in L4, *A*_L4_, at different times after the onset of a brief stimulation to D2 whisker. Bottom: Analogous plots for L6 populations, where population spikes occur later and over longer periods. (B) Temporal profiles of mean firing rate of three cortical columns in L4 (top) and L6 (bottom) respectively. Stimulus duration is indicated in grey shading. The timescale of the left panels matches that in (A) to show the behaviour of the initial population spikes evoked by sensory stimulation. Late spike-bursts in L4 driven by secondary thalamocortical input are displayed over a longer timescale (top-right), but the additional late responses can hardly spread into L6 (bottom-right). (C) Population activity averaged over all 100 TC neurons in D2 barreloid, *A*_TC_. The early synchronous activity is induced by the principal whisker deflection and relays the sensory information to the L4 barrels (left). The trapezoid form of stimulus is illustrated under the plot. The initial L6 population spike (bottom-left panel in (B)) projects back to the thalamus, eliciting the late oscillatory rhythm that begins after ~110 ms of stimulation onset at an interval of ~110 ms (right). (D) Spike raster for the full thalamic network of D2 barreloid (from which the population activity of TC cells shown in (C) is calculated). The circuit is composed of 100 TC cells (dark blue) in the D2 barreloid and 100 reciprocally coupled RE cells (light blue). The time axes of plots in (B), (C) and (D) are aligned.

In addition, a grid of barreloids in the VPM of the thalamus was simulated to convey sensory signals from individual whiskers to the cortex. The activity evoked by the principal whisker in barreloid D2 will not spread into other barreloids because no recurrent connections are made within the VPM, as observed experimentally [51]. The following describes the information flow within the thalamocortical system in response to a D2 whisker deflection: The D2 whisker stimulation first excites the D2 barreloid through a bottom-up connection to TC neurons, and subsequently induces transient PSs in the cortical L4 populations via thalamocortical synapses. Onset PSs are also evoked in L6 through intra-cortical connections and projected back to the thalamus, where the spindle oscillations are initiated after the instantaneous onset response. As can be seen in Fig 2C and D, the TC cell group in the D2 barreloid exhibits a transient increase in population activity right after peripheral stimulation (early response), followed by substantial synchronized activities in the sleep spindle frequency range (roughly 9 Hz, late response). The diminishing late oscillation triggered by corticothalamic excitation occurs about 110 milliseconds after stimulus onset and lasts a couple of cycles before termination. Finally, the late thalamic activity gives rise to a secondary cortical response in L4 that only barely propagates to L6 (Fig 2B, right panels).

### SSA arises from adapted PS to standard stimulation

SSA is generally tested in the oddball protocol, where repetitive deflection is applied to one whisker (called the standard), and the sequence is randomly interrupted from time to time by deflection of another whisker (called the deviant). The roles of whiskers as standard and deviant are swapped to disambiguate any response preference for individual whisker from the deflection probability on SSA [8].

In accordance with experimental findings [8], the L4 network exhibits prominent SSA in the late response but hardly in the initial population spike, where a similar amplitude of activity is elicited by whisker stimulation, regardless of its identity as either standard or deviant (Fig 3A, middle panel). The initial PS is relatively unaffected because the depleted synaptic resources in the L4 populations can always recover almost fully before the next stimulus is presented (Fig 3A, bottom). In contrast, depleted synaptic resources in L6 of the standard column (D2) recover only partially during the inter-stimulus interval, due to the relatively long synaptic recovery time constant in this layer (Fig 3B, blue trace in bottom panel). The insufficient resources, along with depressed L4-to-L6 input in D2 caused by repeated standard stimuli, can only trigger inadequate PSs (Fig 3B, blue trace in top panel). However, as a result of the low probability of deviant stimuli, synaptic resources in L6 of the C2 cluster often reach their steady-state (Fig 3B, red trace in bottom panel) and accordingly evoke full PSs when the deviant stimulus finally arrives (Fig 3B, red trace in top panel). In other words, SSA is initiated in L6 during the early PS phase, where the cortical L6 drive on the deviant barreloid is always strong enough to elicit spindle oscillation, while it is in most cases too weak to do the same on the standard barreloid (Fig 3C and 3D). These late deviant-selective oscillations in the thalamus are then fed back to L4 populations and result in the late L4 rhythmic dynamics exhibiting SSA, consistent with the experimental results that the late cortical responses are very likely to be evoked by secondary thalamocortical inputs to L4 [8].

**Fig 3.**
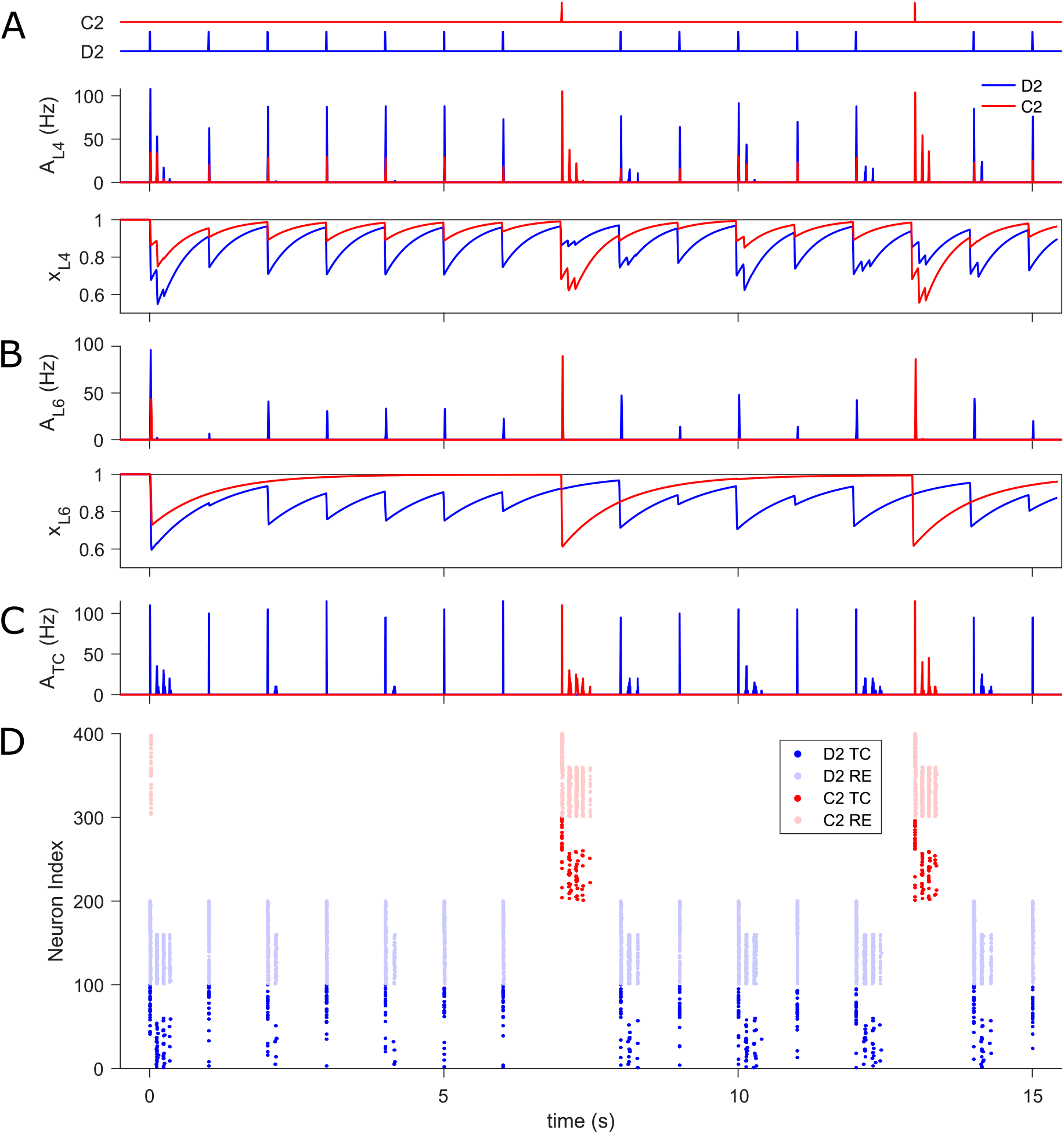
SSA in the thalamocortical network. (A) Time course of the whisker oddball protocol (top), L4 population activity, *A*_L4_, (middle), and mean synaptic resources, *x*_L4_, (bottom) in standard D2 (blue traces) and deviant C2 (red) barrel. Both standard and deviant stimuli evoke comparable amplitude of PS (the first burst in response to each stimulus) due to the fast recovery of depleted resources. Conversely, late oscillations occur primarily in response to deviant stimuli. (B) L6 population activity, *A*_L6_, (top) and mean synaptic resources, *x*_L6_, (bottom) in D2 and C2 infrabarrel. The deviant stimulus generally triggers more substantial PSs than the standard because of more resource available upon presentation of the deviant than the standard. (C) Population activities of TC cells, *A*_TC_, in D2 and C2 barreloids. The deviant corticothalamic PS is always strong enough to initiate late thalamic oscillation, yet this is not the case for the standard. (D) Spike raster of thalamic neurons in D2 (blue) and C2 (red) barreloids. The transient activity of C2 RE cells in response to the first stimulation are caused by the significant cortical cross-column L6 feedback. The time axes of all plots are aligned. In the oddball paradigm, peripheral stimulus duration is 10 ms and inter-stimulus interval (ISI) is 1 s (onset-to-onset).

It is worth noting that late thalamic and synchronized cortical oscillations are also sometimes elicited in D2 by the standard stimulation, and especially the standard stimulus immediately following a deviant (Fig 3C, D and middle panel of A), in accordance with the sporadic standard-triggered oscillation recorded in L4 of the cortex [8]. Here these late rhythmic responses occur largely because the deviant did not deplete as many resources in L6 of D2 as another standard would have done, which occasionally allows strong enough cortical drive on the thalamus to generate a spindle oscillation during the presentation of the next standard stimulation.

### Context-dependent deviance detection

It has been suggested that the larger responses to the stimuli features, when deviant than when standard, not only reflect the rarity of the deviant causing less use-dependent adaptation [3], but also the sensitivity to the violation of the expectation established by the repeated standard, indicating a dynamic prediction mechanism for the next stimulus based on the short-term memory of the previous statistical sensory stream [52]. Such sensitivity to contextual information is termed true deviance detection/sensitivity. A common test for true deviance detection uses many-standards control protocol, where besides the standard-deviant stimuli pair in the oddball sequence, many other different stimuli are presented as the standards [20]. In this control, each of these distinct stimuli occurs with the same probability as the deviant stimulus, which consequently eliminates any potential expectation for the next stimulus but preserves the rarity of the deviant. If the response to the deviant stimulus in the oddball sequence exceeds the response to the same deviant one embedded in the control sequence, then the existence of true deviance detection is confirmed [20]. Here in the many-standards condition for the somatosensory domain, the standard stimuli are equiprobably distributed over three whiskers in a row that are adjacent to the deviant whisker, therefore each whisker is deflected with 25% probability [8] (Fig 4A).

**Fig 4.**
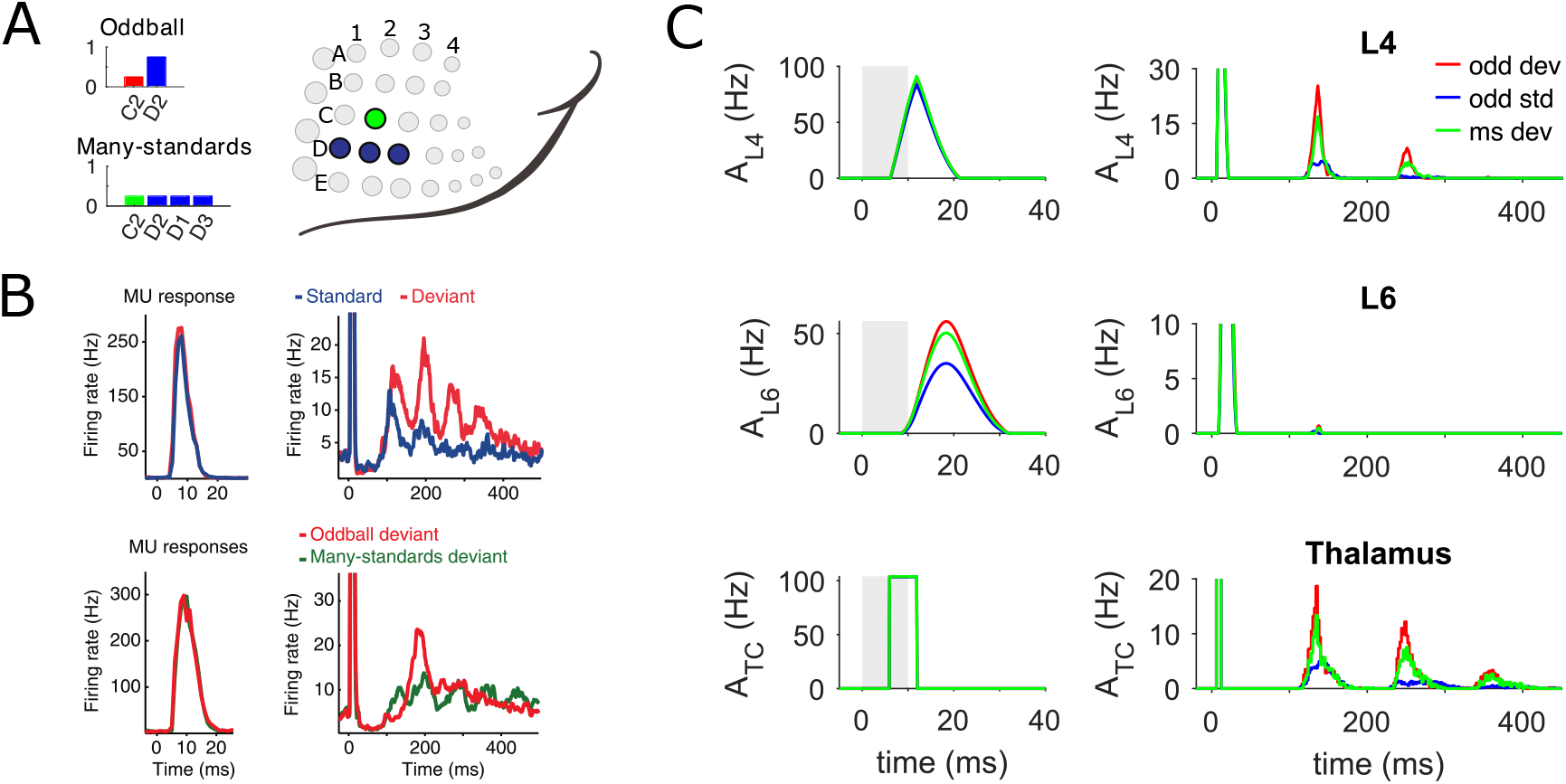
Sensitivity to deviance in silico vs in vivo. (A) Two protocols are used to test context-dependent deviance detection. In the whisker oddball condition, deviant stimuli are applied to the C2 whisker with 25% probability of occurrence, while standard stimuli are applied to the D2 whisker with 75% probability. In the many-standards condition, the deviant stimulus is presented with the same probability of appearance as in the oddball condition, but standard stimuli are equiprobably distributed over three whiskers D1-D3, each of which is stimulated 25% of the time. The bar height represents the probability of a specific whisker being stimulated. (B) A subset of multi-unit (MU) recordings in the granular layer of rat S1 shows late deviant-specific responses in the whisker oddball and many-standards protocols. Average early spike (top row, left) and late deviant-specific oscillatory responses (top row, right) are shown for standard (blue curves, activity recorded in D2) and deviant (red, activity recorded in C2) stimuli in oddball condition. The late response (bottom row) also demonstrates a distinct difference between two deviant types (both responses recorded in C2) in oddball (red curves) and many-standards protocols (green) respectively. Data are adapted from [8]. (C) Average biphasic population activity in cortical L4 (top row), L6 (middle) and thalamus (bottom) in response to oddball standard (blue traces, recorded in D2), oddball deviant (red, recorded in C2) and many-standards deviant (green, recorded in D2) stimuli. Similar to (B), the left and right columns of panels show the early and late activities respectively across cortical layers and the thalamus. SSA and true deviance detection are initiated in the early L6 responses and subsequently enhanced in the late thalamic oscillation, which finally induces the secondary deviant-selective cortical response in L4. The third cycle of thalamic activity is too weak to produce a response in L4 since this thalamic input is below the rheobase (current threshold of gain function) of L4 neurons. Stimulation periods are marked in grey shading. Data in each protocol are averages of 120 stimulus presentations. In both stimulation paradigms, peripheral stimulus duration is 10 ms and inter-stimulus interval (ISI) is 1 s.

Fig 4C illustrates the distinctive biphasic activity of C2 populations evoked by the oddball and many-standards deviant as well as of D2 populations by the oddball standard whisker deflection in the cortical L4, L6 and the thalamus. For comparison, examples of cortical multi-unit (MU) recordings exhibiting late deviant-selective responses in both protocols are replotted in Fig 4B from [8]. The late responses are only found in a subset of recordings mainly confined to the L4. In line with the experimental findings, the simulated L4 populations elicit nearly equally strong population spikes at short latency to both the oddball standard and deviant, as well as the many-standards deviant stimulation, while SSA and true deviance detection are only observed in the late oscillatory responses (Fig 4C, top row). Importantly, our model makes further predictions of the population activity in the cortical L6 and the thalamocortical populations in both somatosensory oddball and control protocols, which, to the best of our knowledge, has not been conducted experimentally yet. We hypothesize that the late context-dependent deviant oscillation seen in L4 is reflected by the thalamic spindle oscillation that can be solely generated by the specific thalamic circuit (Fig 4C, right panel of bottom row). The differences in the secondary thalamic responses to each type of stimulus are driven by the early cortical PS variations in the L6 circuit, which independently detects the deviant in its own responses (Fig 4C, left panel of middle row).

To understand why the average early response of the L6 network for the oddball deviant is larger than for the many-standards deviant, we compared the dynamics of deviant responses in L6 across the two conditions. In the regular oddball sequence, the inadequate L6 activity in D2 evoked by the repetitive standard stimulus hardly propagates into C2, leading to negligible L6 resource depletion in C2 and little cross-whisker adaptation (refer to Fig 3B, bottom). In contrast, in the many-standards sequence, the standard stimuli generate substantial activity in their corresponding clusters that provide C2 with strong inter-column inputs, because of the equiprobable distribution of the standard stimuli and accordingly longer recovery time of resources between repeated stimulation. This additional lateral input generally results in higher activity in C2 evoked by the standard stimulation in the many-standards condition than the oddball condition, and consequently leads to more adaptation. To illustrate this mechanism for true deviance detection across the hierarchical network, Fig 5A–5E displays population activities and synaptic resources for the C2 cluster in both protocols across the L4, L6, and thalamocortical populations respectively. The dashed brown frames highlight two examples for an oddball deviant triggering a significant late response in L4, while a many-standards deviant does not. In the first frame, the thalamocortical input as a deviant in both protocols triggers approximately the same onset PS in L4 of C2 (Fig 5B, top, first spike within the box). In contrast, less L6 activity is elicited in the many-standards than the oddball condition owing to the lower level of L6 resources reduced by the last D3 stimulation in the many-standards condition (Fig 5C). In consequence, only the corticothalamic feedback in the oddball protocol is large enough to trigger spindle oscillations in TC cells (Fig 5E), which in turn cause synchronized activity in the cortical L4 barrel (Fig 5B, top). In a similar vein, the second frame shows an example where resources in L6 of C2 in the many-standards condition are freshly exhausted by the preceding D2 stimulation and do not have time to recover adequately to trigger a spindle oscillation. It is also worth noting that under cross-whisker adaptation, the resources in L6 of C2 are occasionally depleted by bursting activity at long latency propagated by the first robust burst of late oscillation evoked in L4 of standard column(s) (Fig 5C, an instance is highlighted by golden pyramid).

**Fig 5.**
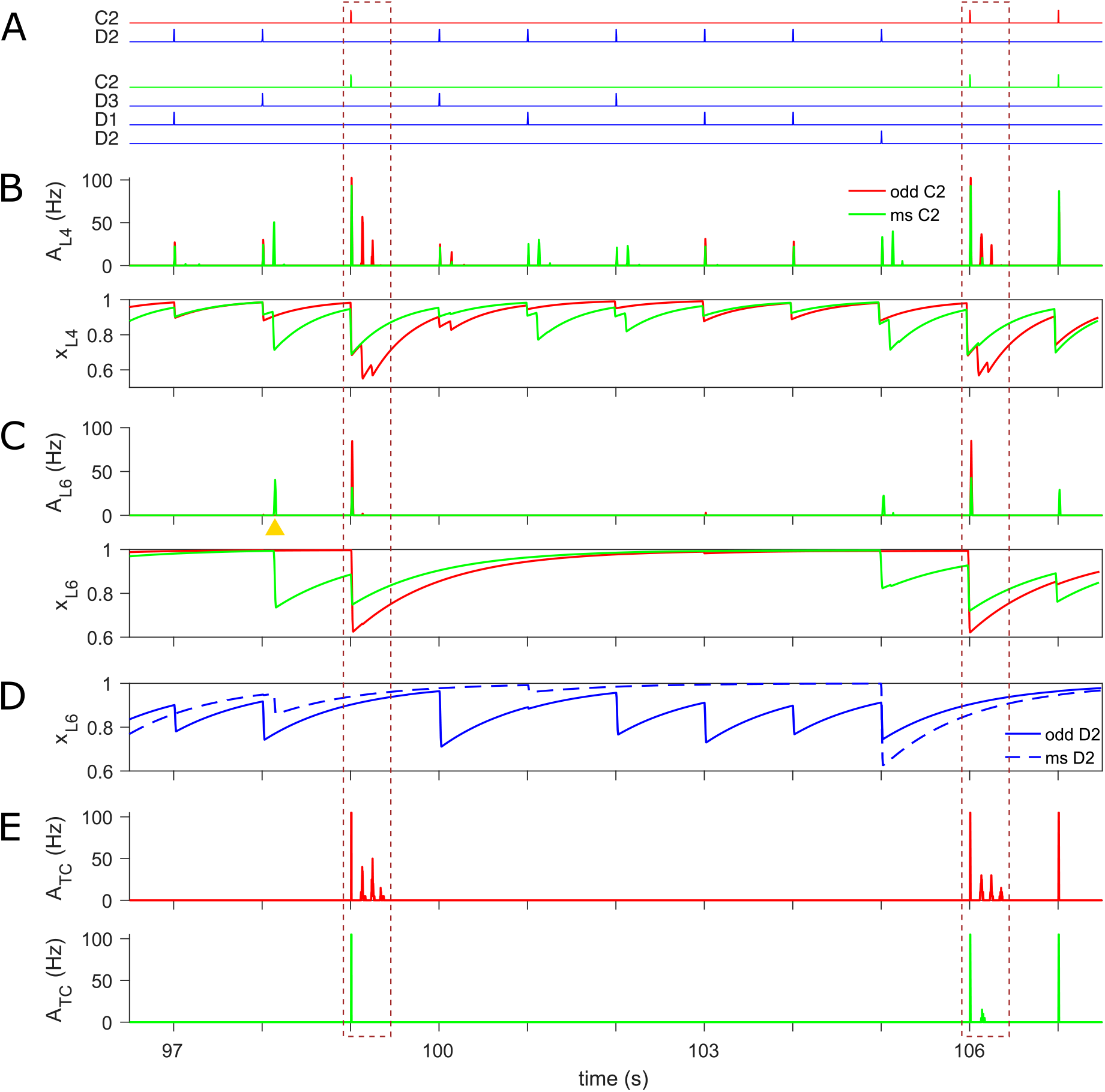
True deviance detection in the model network. (A) Illustration of oddball (top) and many-standards (bottom) protocols. The deviant stimulation to C2 is highlighted in red for the oddball condition and in green for the many-standards one. (B) Population activity (top) and synaptic resources (bottom) in L4 of C2 in response to deviant stimuli presented in oddball (red traces) and many-standards (green) condition respectively. The deviant in both protocols often evokes comparable onset responses, while a robust late oscillation is generally only induced by the oddball deviant (two typical cases are highlighted in dashed boxes). (C) Same as (B) but displayed for L6 of C2. Considerably larger early onset responses to the many-standards deviant compared to the oddball deviant are generated in the L6 population (see dashed boxes), because higher activity is propagated from the L6 standard column(s) into that of the deviant column in the many-standards compared to the oddball condition, and accordingly less synaptic resource is left and a weaker response is evoked upon arrival of the deviant. (D) Synaptic resources in L6 of D2 in response to the standard stimulus applied to D2 whisker embedded in oddball (solid traces) and many-standards (dashed) sequences. The standard stimulus to D2 in the many-standards condition, with its lower probability of appearance than the same stimulus in the oddball condition, allows the resource in L6 of D2 to have more time to recover fully before the next standard stimulus arrives and consequently to generate an adequate PS capable of spreading into the same layer of the deviant column. (E) Thalamocortical (TC) population activity in C2 barreloid evoked by the deviant in the oddball (red) and many-standards (green) protocols. The corticothalamic excitation of the deviant in the many-standards condition is normally strong enough to elicit late intermittent bursting, whereas that of the deviant in the oddball condition is not (dashed boxes). The time axes are aligned for panels A-E. In both stimulation paradigms, peripheral stimulus duration is 10 ms and inter-stimulus interval (ISI) is 1 s (onset-to-onset).

### Further experimental predictions

We ran another pair of novel prediction paradigms on our model with the same set of parameters used in all previous protocols to test the model’s sensitivity to contextual information that includes sequential statistics. Here we used variants of the many-standards paradigm with fixed deviant position but permutated the order of the standards presentations to be either random or periodic. In the sequenced paradigm, the standard deflection was periodically applied to C2, D1 and B1 whiskers, each of which is rarely substituted for the C1 deviant in order, which is overall presented 3% of the time. Identical statistics were used in the randomized paradigm except that the three standard stimulations are randomized between the successive deviants. A schematic representation of both sequences is displayed in Fig 6A.

**Fig 6.**
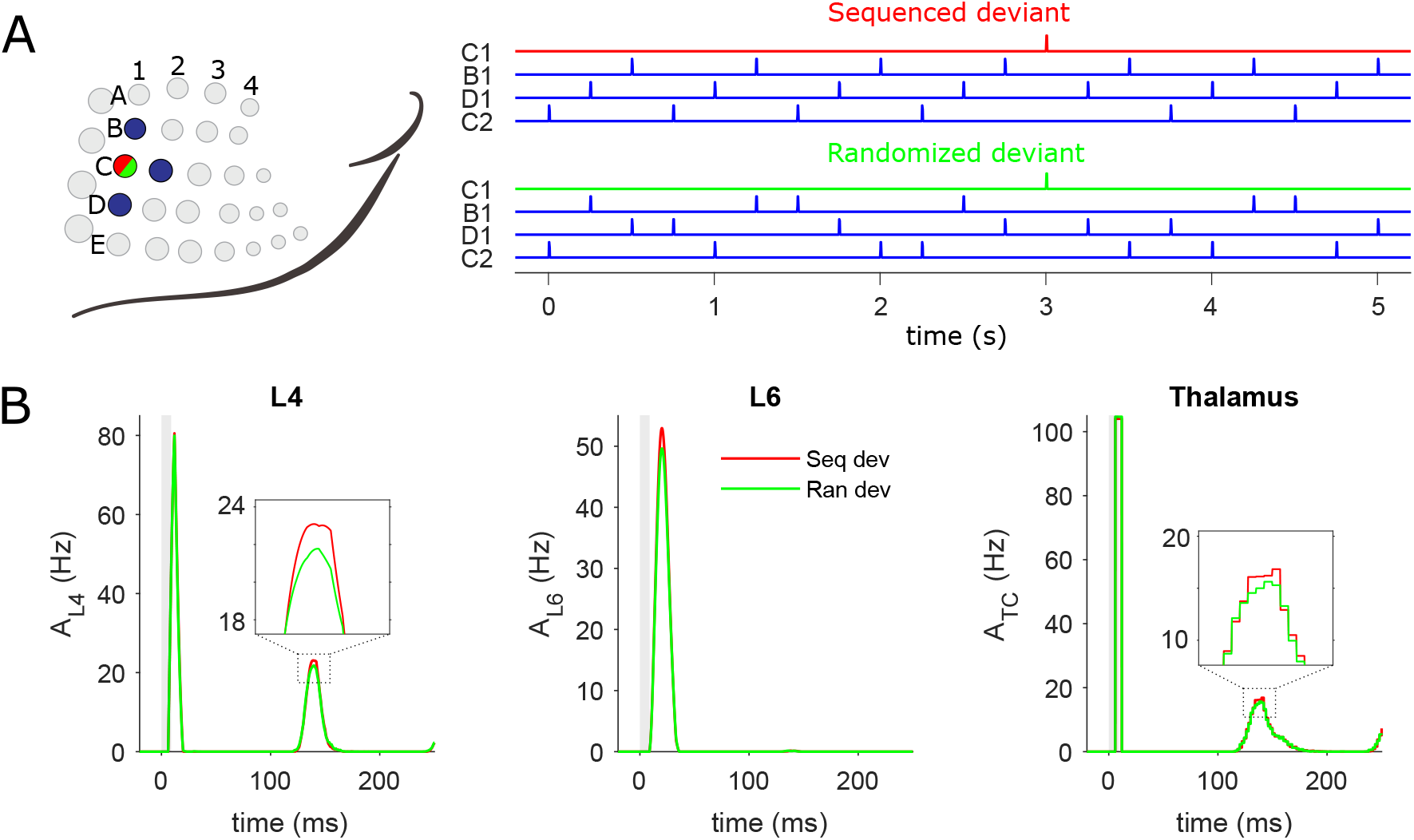
Deviant responses of cortical and thalamic populations in sequenced and randomized paradigms. (A) Schematic illustration of sequenced and randomized stimulation trains and the stimulated whiskers. In two sequences, the delivery of the deviant stimulus to the C1 whisker is the same, but three standard stimuli are employed either periodically (sequenced condition, top) or randomly (randomized condition, down) to C2, D1 and B1 whiskers. In the sequenced stimulus train, the deviant occasionally breaks the regularity by repeatedly taking the place of one of the standards in the order of C2, D1 and B1. In both paradigms, stimulus duration = 10 ms and inter-stimulus interval (ISI) = 250 ms (stimulus offset to onset). Deviant stimuli constitute 3% of overall stimuli (30 out of 1000 stimuli). (B) Average population activity across cortical L4 (left), L6 (middle) and thalamus (right) in response to the deviant in sequenced (red) and randomized (green) paradigms. A weakly differential response to sequenced and randomized deviant exclusively appears in the L6 population at short latency. Owing to the thalamocortical loop, this subtle difference is successively manifested in late thalamic and cortical L4 bursting activities, which are magnified in insets for clarity. Stimulus duration is highlighted in grey. CSI values are 0.0115, 0.0254 and −0.0061 for early L6 (computed over [0, 40] ms) and late L4 and thalamic responses (computed over [40, 250] ms), respectively.

In comparison with the sequenced condition where the expectation is established by the repeated sequential standards and violated by the deviant, a precise prediction about forthcoming stimuli cannot be formed in the randomized condition. As shown in Fig 6B, a slightly stronger L6 response is initiated for the deviant embedded in a sequenced, expectation-forming than random stimulation train. The response discrepancy is also reflected in L4 with latency by the thalamocortical loop. We quantify this effect by computing the context-specificity index (CSI) defined as the difference between responses to sequenced and randomized deviants normalized by the sum of two responses.

Furthermore, our model predicts the dependence of the novelty-predicting effect on the inter-stimulus interval (ISI) across different cortical layers and thalamus (Fig 7). Due to the feature of biphasic dynamics in our network, CSI is separately computed for the early and late responses. For the early response, the novelty-predicting effect is demonstrated in the cortex over a wide range of ISI and particularly prominent in L6 that as we expected to be the novelty detector. Early novelty-predicting responses originated in L6 are subsequently revealed in the thalamic oscillation and secondary L4 responses via the L6-to-thalamus-to-L4 pathway.

**Fig 7.**
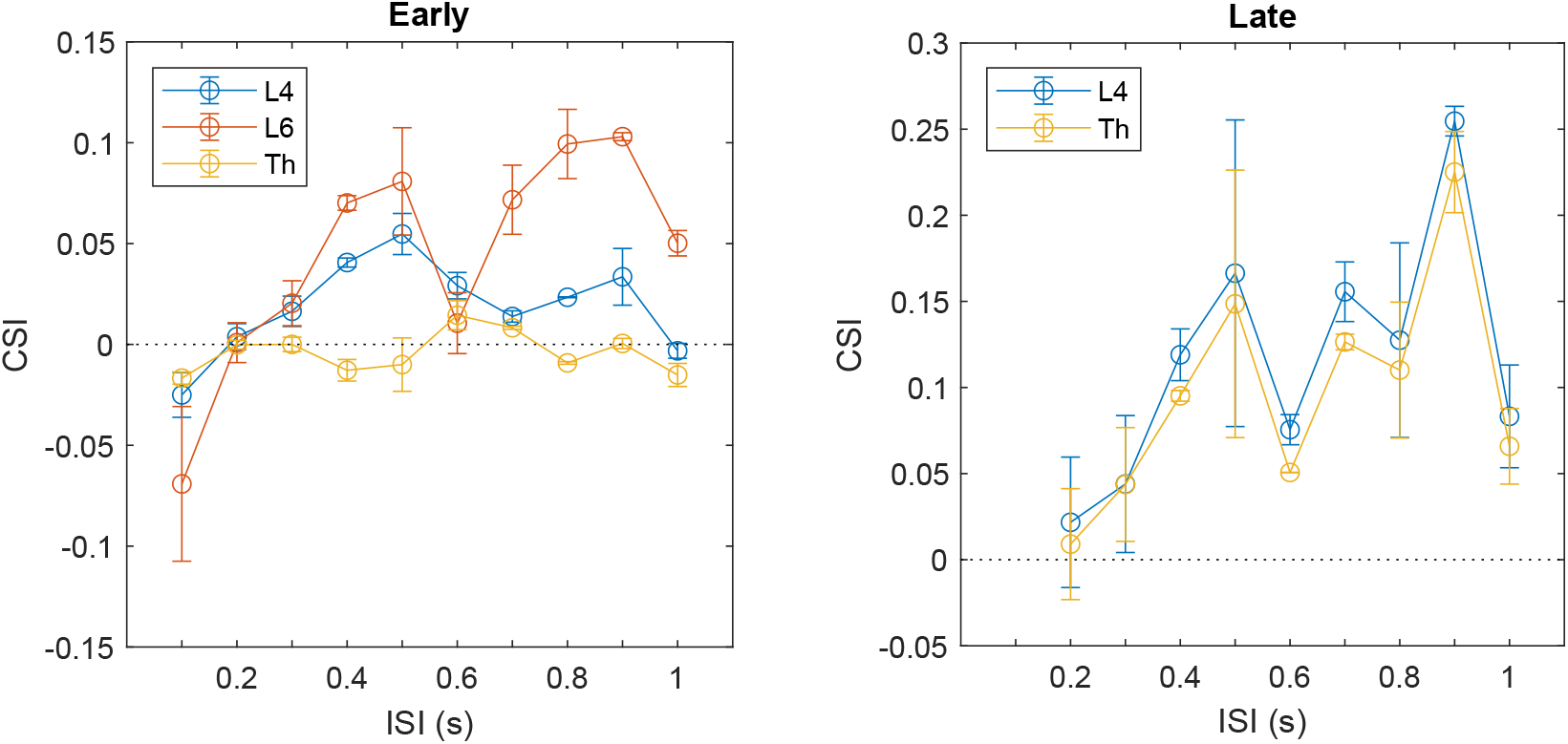
Novelty-predicting effect as a function of inter-stimulus interval. Early and late CSIs are respectively computed over 40 ms after stimulus onset and the remaining time before the onset of the next stimulus. Data are averaged over 5 trials of each paradigm consisting of 1000 stimuli.

To eliminate the potential effect of sensory input uncertainties on CSI, we removed the stimulus-by-stimulus variation of the thalamocortical drive that was previously presented and reran both protocols on the model. In this case, each external stimulus evokes the same amplitude of thalamic responses. One can clearly see from Fig 8A that the novelty-predicting effect is greatly attenuated near both ends of the range allowing for it because either of the cases will lead to the steady-state of unrecovered/recovered synaptic resources in L6, which are independent of the presentation context. We also investigate the individual role of three types of deviant in the sequenced paradigm in detecting novelty across different ISIs, where the C1 deviant preceded by the B1, C2 and D1 standard is defined as type 1, 2 and 3, respectively (Fig 8B). The novelty-predicting effect is the most pronounced for the type 2 deviant and is not influenced much by the increase in ISI after the first several values, which indicates the capacity of novelty prediction in the cortex depends on the specific sequential context in the recent past in randomized paradigms, i.e., the precise information about the order of standard presentations over hundreds of milliseconds before the arrival of deviant.

**Fig 8.**
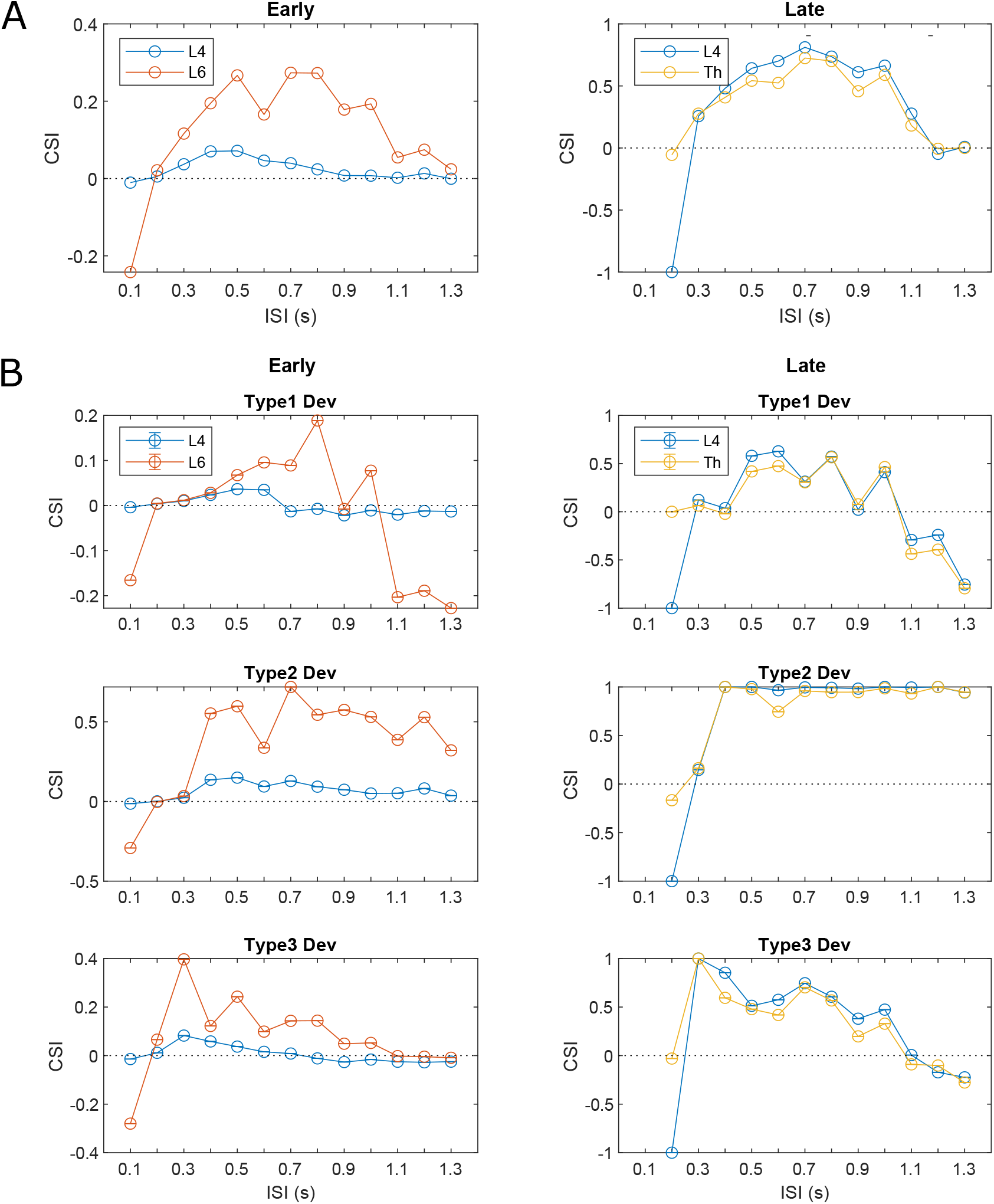
Novelty-predicting effect as a function of inter-stimulus interval in the model with deterministic sensory inputs. (A) Early and late CSIs are computed in the same fashion as Fig 8 except that data are averaged over 30 deviant out of 1000 stimuli in each paradigm. The CSI values for the early thalamic responses are ignored since they are always zero independent of ISI. (B) Three types of deviants show different dependence of novelty predicting capacity on ISI. Type 2 deviant has a wider ISI range allowing novelty prediction in both early and late responses than the other two types.

## Discussion

We presented a cortico-thalamo-cortical circuit of somatosensory processing with recurrent dynamics and depressing synapses that is capable of producing both SSA and the contextual sensitivity over different latencies across cortical layers and thalamus, in line with previous experimental observations [8]. The computational results suggest that the early context-specific deviant responses in S1 L6 could be a strong candidate for the electrophysiological correlates of the mid-latency response observed in EEG recordings [22, 26], because of their similar response latencies and shared sensitivity to presentation context. Besides, we show that the late context-dependent responses in S1 L4, which temporally and functionally resemble MMN, can be produced simply by the interaction between different cortical layers and the thalamus, without involving top-down modulation by higher brain areas.

Inspired by a previously described mechanism producing SSA through interactions among intracortical populations with different specificity for stimulus features in the auditory cortex [35], our model enables the generation of a cortical population spike upon the presentation of stimuli and its lateral propagation across different barrels to mediate SSA, consistent with the given neurobiology of the barrel cortex. The existence of whisker-evoked population spikes and their spatiotemporal distribution in S1 are inferred from voltage-sensitive dye imaging [53] and multielectrode array recordings [54, 55], where the mean firing rates show a temporary peak in the principal column upon the onset of stimulation, followed by weaker activity with short latency in the surrounding columns [55]. Previous modelling work has demonstrated that this population burst could be a consequence of a recurrent network with synaptic depression [43, 44, 56, 57]. Here, such a network is the essential ingredient of our hierarchical model to produce SSA.

Our model can be regarded as a simplified representation of the cortico-thalamo-cortical pathway of the rodent whisker system. The dynamics of the barrel cortex are described by population activity and mean synaptic resources of somatotopic neuronal clusters in a two-layered grid network. The reasons for adopting rate models for the cortical dynmaics are twofold. First, the mean-field description of population activity makes it possible to analyze the dynamics of single recurrent clusters giving rise to population spikes on the phase plane, under the mathematical abstraction that each cortical cluster is a large and homogenous network. Theoretical work has demonstrated that such a mean-field representation is equivalent to a homogeneous pool of leaky integrate-and-fire neurons in terms of its population activities [58]. Second, this specification allows us to focus on the collective population behaviour within the grid network, which is sufficient to generate cortical SSA and true deviance detection. SSA and true deviance sensitivity cannot be accounted for solely by the intrinsic dynamics of individual neurons, but relies to a large extent on network dynamics. It should be noted that although this mean-field network of the barrel cortex ignores some details, such as the potential contribution of additional layers, distinct neuron types and their morphology, and more realistic connectivity profiles, critical features of the barrel cortex including its somatotopic columnar arrangement and the spatiotemporal pattern of whisker-evoked responses are retained and indeed shown to be sufficient to explain the deviant selectivity of transient L6 responses.

For the thalamic contribution, following previous work by Destexhe et al. [46], a spiking neural circuit of single-compartment thalamocortical (TC) and thalamic reticular (RE) cells are used to reproduce spindle oscillation. Synaptic currents within the network are simulated by the synaptic kinetics of AMPA and GABA receptors. In contrast to the cortical rate model, which only depends on network mechanisms, the thalamic spiking model takes into account both the intrinsic properties of single cells and their interplay to elicit synchronous oscillations. Specifically, the spiking dynamics of each thalamic neuron is simulated by an Izhikevich model that reproduces similar firing patterns to the conductance-based model used in [46], but is computationally as efficient as the leaky integrate-and-fire model. Our modelling results implicate the slow recovery variable in these thalamocortical neurons, which may describe the inactivation of low-threshold Ca^2+^ T-currents [59], as the main factor driving the latency of the secondary response observed in S1 L4 and therefore potentially MMN.

Finally, it is worth mentioning that the current implementation of our model still has several shortcomings. A major limitation is the sensitivity of SSA to cortical network parameters, which stems from the different layer-specific response patterns in the feedforward L4-to-L6 network. In other words, the parameter regime allowing for SSA and the true deviance sensitivity in initial L6 responses is constrained by not only the spatiotemporal pattern of whisker-evoked transient activity in the L6 network but also responses in L4 that directly delivers its output to L6. Primarily, in the L6 circuit, the PS evoked by the oddball standard should reduce with repeated stimuli while not generalize to the oddball deviant. Besides, the propagation of activity across the L6 network is supposed to be robust enough to occasionally pass the PSs triggered by standards in the many-standards condition into the deviant column, yet too weak for PSs evoked by the oddball standard to do so. Another possible shortcoming of the model is that it omitted cortical layer 2/3 and layer 5 that could modulate the laminar pathway from L4 to L6. Future work incorporating these missing layers could generate richer dynamics that expand the narrow region of SSA existence in the parameter space, leading to a robust SSA phenomenon. Lastly, it would be instructive to test the NMDA dependency of the late response in a spiking implementation of the cortical dynamics explored here to further investigate SSA’s connection with MMN.

## Materials and methods

### Derivation of spiking to rate network

We derive the rate equations for a network of interacting populations mediated by depressing synapses assuming that each population is large and homogeneous, using standard techniques [58, 60–62]. By homogenous, we mean that (i) all neurons have the same properties; (ii) each neuron receives identical external input; and (iii) the coupling strength between any pair of neurons within the population is statistically uniform. The general conclusions developed here will be applied in the next section to formalize the mean-field cortical network composed of a grid arrangement of coupled populations.

We define the mean firing rate as the population activity *A*(*t*) averaged over a pool of homogenous neurons.

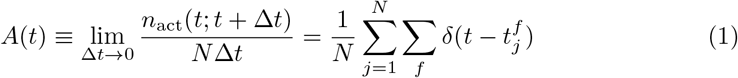

where *N* is the number of neurons within the population, *n*_act_(*t*; *t* + Δ*t*) denotes the number of spikes of all neurons in the population, which appear betweem time *t* and *t* + Δ*t*. Since Δ*t* is a very short time interval, each neuron fires at most one spike during the interval and correspondingly *n*_act_(*t*; *t* + Δ*t*) also represents the number of neurons that fire between time *t* and *t* + Δ*t*. *δ*(*t*) is the Dirac delta function and 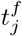 denotes the firing time of the *j*-th neuron in the population. This notion of the rate as population activity averaged over many neurons will be widely used in the following derivation.

The dynamics of short-term depression for the connection between pre- and post-synaptic neuron pair *j*, *i* is given by the phenomenological model [56, 63]:

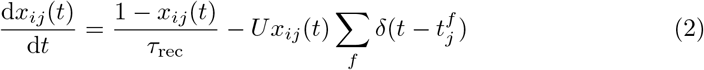

where each pre-synaptic spike emitted at 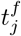 utilizes a certain rate *U* of the available synaptic resource *x_ij_*that is a fraction of full resources. The depleted available resource then returns to its baseline value with a time constant *τ*_rec_. For the current-based type of post-synaptic input, which is independent of neuronal membrane potential, the increment in the amplitude of post-synaptic current *I_i_*(*t*) triggered by a pre-synaptic spike arriving at 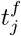 is given by

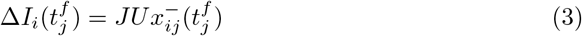

where *J* is the absolute synaptic efficacy and 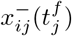 denotes the value of *x_ij_* immediately before the arrival of the spike.

We define 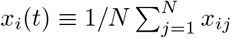 as the mean available resource in a bunch of synapses from a population of pre-synaptic neurons *j* to a post-synaptic neuron *i*. By taking the average over the pool of pre-synaptic neurons on both sides of Eq 2, we get

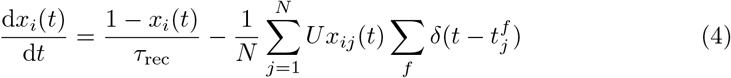

The second term on the right-hand side of Eq 4 can be further transformed, assuming uncorrelated Poisson spiking from a large population of pre-synaptic neurons, to

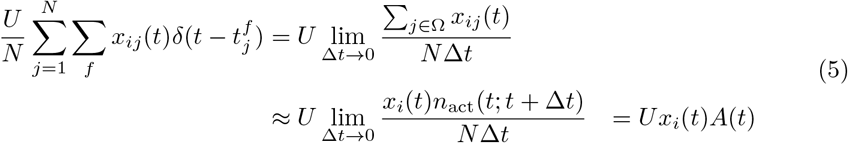

where *N* is the number of neurons in the pre-synaptic population. Ω denotes the subpopulation of pre-synaptic neurons that fire between time *t* and *t* + Δ*t*, and *n*_act_(*t*; *t* + Δ*t*) is the size of the subpopulation. The approximation becomes an equality in the limit of *N* → ∞. *A*(*t*) represents the population activity of pre-synaptic neurons as defined in Eq 1.

Using Eq 5, Eq 4 can be reformulated as

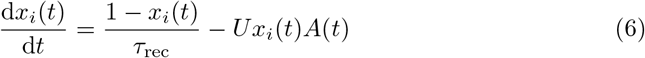

We find that the dynamics of the mean synaptic resource *x_i_*(*t*) is independent of the neuronal identity and therefore is the same for all neurons in the population.

In addition, we assume that each pre-synaptic spike emitted at 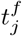 triggers an instantaneous post-synaptic current with the temporal profile 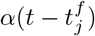 and the efficacy for each synaptic connection in an all-to-all coupled population of *N* neurons is scaled as *J*_0_/*N*. The input current to a neuron *i* in the population is generated by all spikes of all neurons within the same population,

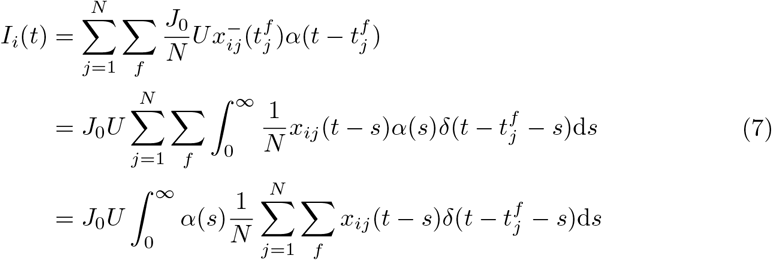

Making use of Eq 5 with t replaced by *t* – *s* to substitute the average quantity on the right-hand side of Eq 7, we obtain

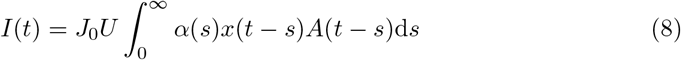

We dropped the neuronal index *i* since the input current at time *t* depends on past mean synaptic resource and population activity and is identical for all neurons.

Now we generalize the arguments from a single fully-connected pool to multiple interacting pools. Under the assumption that neurons are homogenous in each population, the activity of neurons in population *k* is

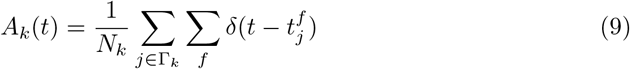

where *N_k_* is the size of population *k* and Γ_*k*_ represents the set of neurons that belongs to population *k*.

We assume that each neuron *i* in population *k* receives input from all neurons *j* in population *l* with adapting coupling strength (*J_kl_*/*N_l_*)*U_kl_x_ij_*(*t*) and the post-synaptic current of the neuron *i* elicted by a spike of a presynaptic neuron *j* has the time course *α_kl_*(*t*). Here the synaptic efficacy *J_kl_*, utilization rate *U_kl_* and post-synaptic current *α_kl_*(*t*) depend on the type of synaptic connection from a neuron in population *k* to a neuron in population *l* rather than the neuronal identity. The input current to a neuron *i* in pool *k* is induced by all spikes of all neurons in the network of pools,

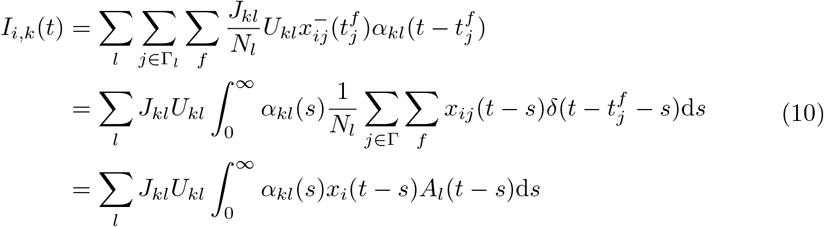

Correspondingly the dynamics of mean synaptic resource *x_i_*(*t*) is generalized as

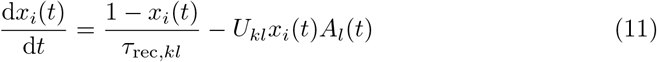

By assuming that *U_kl_* and *τ*_rec,*kl*_ are homogenous across different pools, we drop the subscript *kl* and reformulate the above equation as

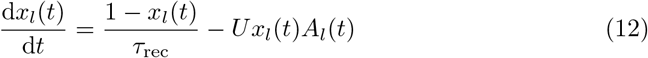

Here we change the index of mean synaptic resource from post-synaptic neuron *i* in population *k* to pre-synaptic population *l* since the evolution of mean synaptic resource is governed by the activity of pre-synaptic population and is the same for all neurons within post-synaptic populations.

In this case, the input current,

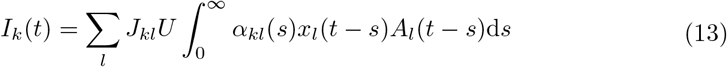

which is independent of the neuronal index *i* but the post-synaptic population index *k*.

Finally, we characterize the dynamics of the input current *h_k_*(*t*) of population *k* with the differential equation of passive membrane and employ for each population the rate model *A_l_*(*t*) = *F_l_*(*h_l_*(*t*)) in which *F_l_* is the stationary gain function of neurons in population *l*. The input current *h_k_*(*t*) takes into account both synaptic coupling *I_k_*(*t*) and external drive 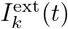,

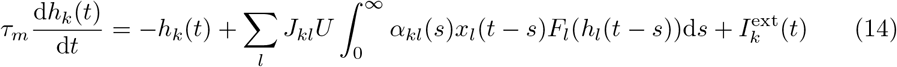

where all neurons in the network have the same membrane time constant *τ_m_*.

Particularly, in the case of instantaneous post-synaptic current pulse *α_kn_*(*s*) = *δ*(*s*), Eq 14 can be reduced to a first-order differential equation

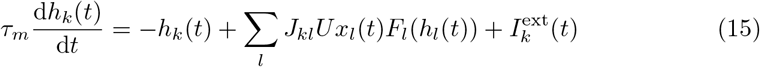

### Modelling the thalamocortical network of the barrel cortex

It has been observed that a cortico-thalamo-cortical loop structure exists in the rodent whisker system. The excitatory thalamocortical (TC) neurons in the ventral posterior medial (VPM) nucleus present feedforward projection to layer 4 (L4) of the cortex. On the other hand, neurons in layer 6 (L6) provide feedback excitation to TC and reticular(RE) neurons that impose reciprocal inhibition on TC cells. Here, for the complexity needed for the work, we greatly simplified the laminar structure of the cortex by only considering two cascaded excitatory networks that model L4 and L6 respectively.

To mimic the somatotopic organization of the barrel cortex, the circuit of each layer is composed of an array of cortical columns (barrels). Each column is modelled by a fully connected excitatory population mediated by synaptic depression, with its mean-field dynamics as analyzed in the last subsection. Every column is also connected to its nearest vertical and horizontal neighbours by inter-column depressing synapses. Furthermore, the L4 columns receive thalamic inputs with cross-whisker tuning and send outputs to their aligned L6 columns.

Based on Eqs 15 and 12, the mean-field dynamics of cortical layer 4 network is described by the following equations (see Fig 9 for a schematic representation of the network architecture):

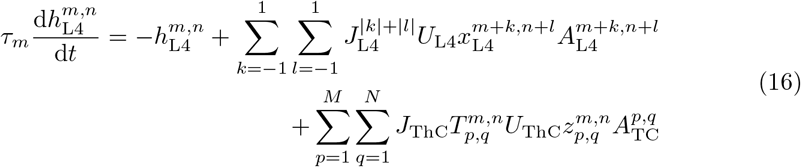

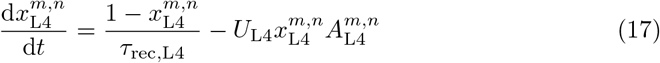

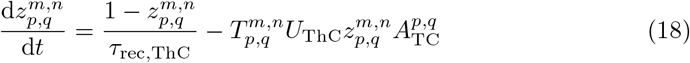

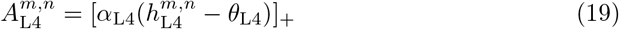

**Fig 9.**
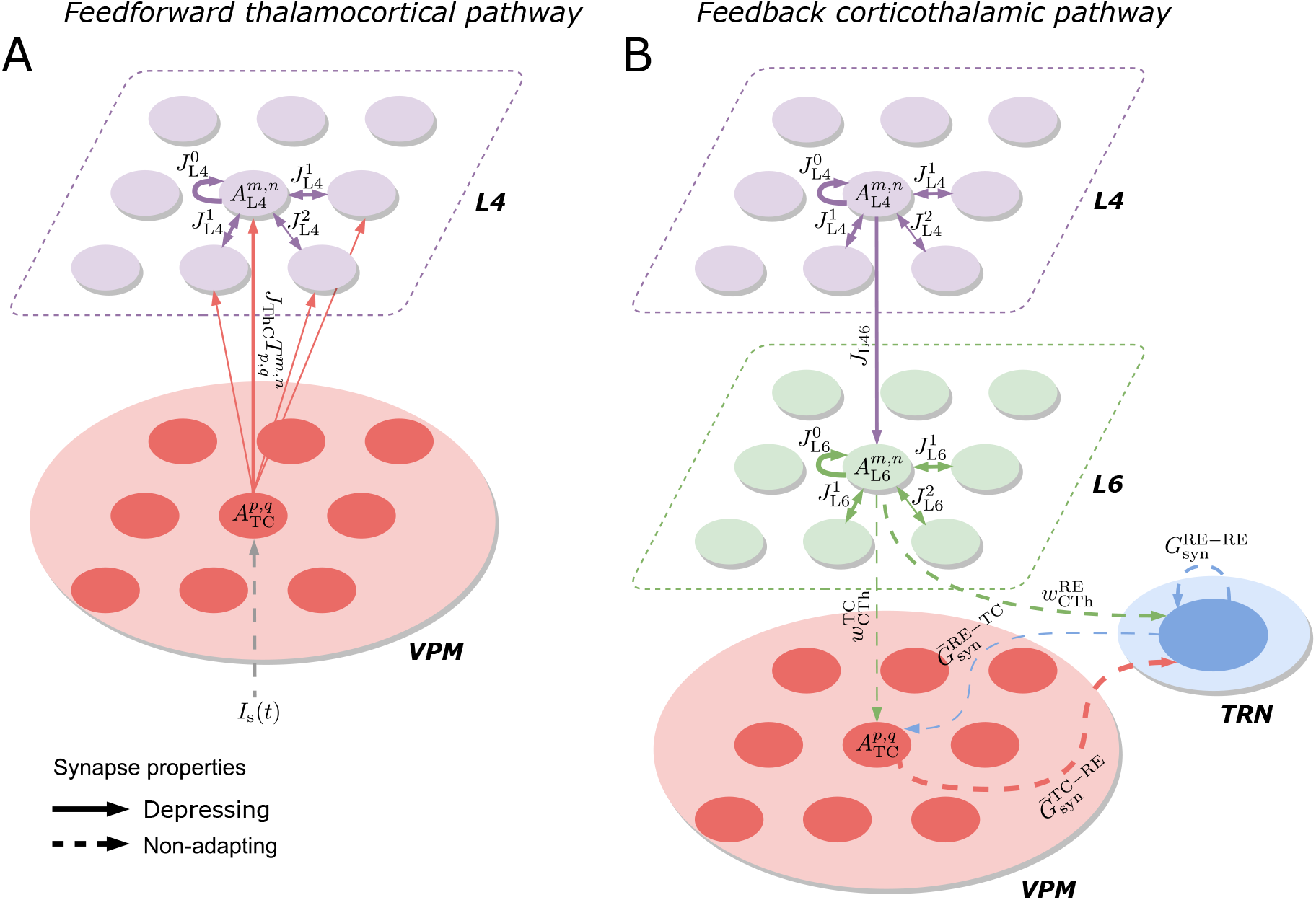
Schematic representation of the thalamo-cortio-thalamic loop model of the barrel cortex, to illustrate the notation used in Methods. (A) Feedforward thalamocortical pathway. (B) Feedback corticothalamic pathway. The width of arrows describes the relative strength of each connection.

Similarly, the mean-field cortical layer 6 network is defined as:

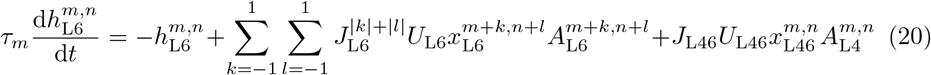

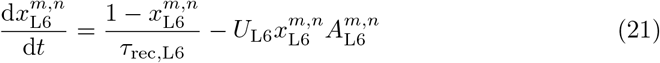

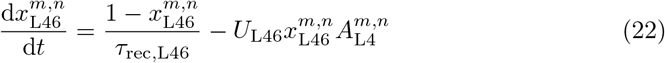

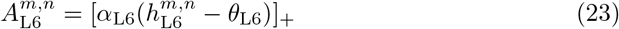

The dynamics of each barrel is described by its population activity *A*^*m*, *n*^ and mean fraction of synaptic resource available *x*^*m*, *n*^, where *m* and *n* denote the row and arc indices of the barrel on an idealized *M* × *N* grid. Both variables are specified by subscript either L4 for the population in layer 4 or by L6 for layer 6 (which applies for all layer-specific variables). The population activity *A*^*m*, *n*^ is defined as a threshold-linear gain function of the mean-field current *h*^*m*, *n*^ (*α* and *θ* are respectively the slope and horizontal intercept of the gain function). The function [·]_+_ is defined as its argument when the argument is positive, otherwise zero.

The synaptic efficacy is represented by *J*^|*k*|+|*l*|^ with superscript specifying the column-to-column distance of the connection (0, 1 and 2 respectively indicate intra-column, vertical/horizontal and diagonal inter-column connection). *U* denotes the utilization rate of synaptic resources. *τ_m_* and *τ_rec_* are the membrane time constant of cortical neurons and recovery time constant of synaptic resources, repectively. The dynamics of thalamocortical synapses from the whisker channel *p*, *q* to the L4 barrel *m*, *n* is characterized by the fraction of resources available 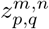, parameterized by utilization rate *U*_ThC_ and recovery time constant *τ*_rec,Thc_. Similarly, the L4-to-L6 depressing synapse of column *m*, *n* is described by its resources 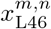 with utilization rate *U*_L46_ and recovery time constant *τ*_rec,L46_. *J*_Thc_ and *J*_L46_ denote the efficacy of thalamocortical and laminar synapses, respectively.

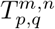 represents the relative magnitude at which the barrel *m*, *n* receives thalamocotical input from whisker channel *p*, *q* compared with its primary whisker channel. The values of 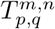 over all channels on the *M* × *N* grid compose the spatial tuning curve of the thalamocortical inputs to the barrel *m*, *n*. We chose a linear tuning curve in the simulation, which is defined as

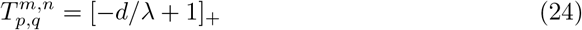

where λ is the radius of tuning curve and *d* is the floored Euclidian distance between barrel *m*, *n* and whisker channel *p*, *q* on the evenly-spaced gird where the separation distance is 1

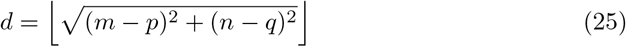

where ⌊·⌋ denotes the floor function that rounds its argument to the nearest integer less than or equal to that argument.

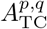 describes the thalamocortical input delivered through the whisker channel *p*, *q*. It is defined as the population activity of TC cells organized in barreloid *m*, *n*, which in practice is evaluated by Eq 1 with a time window Δ*t* = 2 ms. It is worth noting that the time bin of 2 ms is only used to compute the population activity of TC cells from their spikes, and the cortical population activity is obtained by numerically solving those differential equations of the cortical dynamics with an integration time step of 0.1 ms. Specifically, the 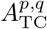 is updated every time window (2 ms) and held constant for 20 integration time steps (each 0.1 ms).

We assume that the late cortical rhythmic activity (roughly 10 Hz) that lies in the frequency range of the spindle oscillation (7-14 Hz) [45] emerges from the thalamo-cortical system. Earlier studies have shown that spindle oscillation is initiated in the thalamus and synchronized over the cortex and its minimal neural substrate lies on the interaction between thalamic RE cells and TC relay cells [45, 46].

Motivated by the anatomy of the VPM and reticular nucleus of the thalamus, we modelled each barreloid in the VPM nucleus by a cluster of 100 TC cells mediated by a pool of 100 RE cells, whose connectivity is illustrated in Fig 9B. The spiking dynamics of each thalamic neuron simulated by the Izhikevich model, which uses the quadratic integrated-and-fire model for the membrane potential equation, is expressed as

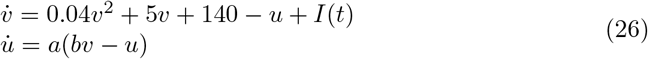

with after-spike resetting

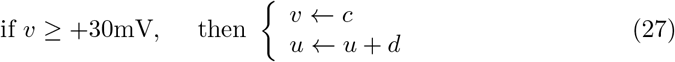

where the time *t* is in milliseconds; *v* and *u* respectively denote the membrane potential (in mV) and recovery variable that represents the difference of all influx and efflux of voltage-gated ionic currents. In particular, the population of 100 TC cells in each barreloid are divided into two subgroups. (*a*, *b*, *c*, *d*) = (0.005, 0.26, −52, 2) are assigned to the first 60 TC neurons (indexed from 1 to 60) to induce rebound bursts that is an essential ingredient of spindle oscillation. The same parameter quartet except for *b* = 0.25 is assigned to the rest 40 TC neurons (indexed from 60 to 100) that only fire spike trains following stimulation from depolarized currents but fail to produce rebound spikes. Thalamic RE cells are modelled as bursting neurons by assigning (*a*, *b*, *c*, *d*) = (0.02,0.2, −55,4). The network connectivity is illustrated in Fig 1C, with a connection probability of 0.6 for TC-to-RE, RE-to-TC and RE-to-RE coupling in each pair of subgroups of TC and RE cells and 0.2 for coupling between two subgroups of RE cells. *I*(*t*) is the total current input to the the neuron (in pA)

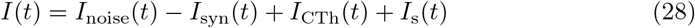

The thalamic background noise *I*_noise_(*t*) is subject to uniform distribution bound between −0.5 and 0. The conductance-based synaptic current is modelled as

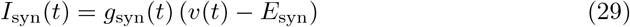

where *E*_syn_ is the synaptic reversal potential. Synaptic conductivity *g*_syn_(*t*) has a time course of exponential decay:

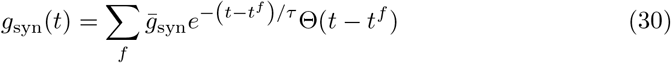

where Θ(*t*) is the Heaviside step function and *t^f^* represent the arrival time of pre-synaptic spikes. We simulated AMPA and GABA_A_ receptors with the decay time constant *τ* = 5 and 6 ms; *E*_syn_ = 0 and −75 mV, respectively. The maximal conductance 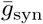 for each type of connection is scaled as 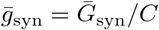, where *C* = 100 × 0.6 = 60 is the number of randomly chosen presynaptic partners for each neuron and 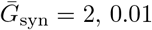 and 0.2 nS for AMPA-mediated TC-to-RE, GABA_A_-mediated RE-to-TC and GABA_A_-mediated RE-to-RE synapses, respectively.

In addition, half of TC cells in each barreloid and their coupled RE cells were randomly picked receiving corticothalamic feedback current, which is expressed as

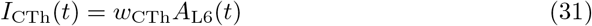

where *A*_L6_(*t*) denotes the population activity of L6 neurons in the somatotopic infrabarrel. We respectively assigned the corticothalamic coupling strength *w*_Cth_ = 0.001 and 0.4 for TC and RE neurons.

Finally, we respectively stimulated randomly 10% and 50% of TC cells in the two complementary subgroups of each barreloid with sensory input that is described as

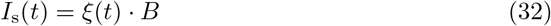

where *B* and *ξ*_*p*,*q*_(*t*) respectively represent the maximum amplitude and temporal envelope of the sensory stimulus (normalized between 0 and 1). In all simulation protocols, *B* = 5 and *ξ*_*p*,*q*_(*t*) has the profile of trapezoid pulse with 10 ms duration (2 ms onset/offset ramp time).

Forward Euler method with a time step of 0.1 ms is used to simulate the network dynamics. The values for different network parameters are listed in table 1. The Matlab code for running the network model is available online at https://github.com/ChaoHan-UoS/SomatosensorySSA_Model.

**Table 1.** Values used for the network parameters

| Notation | Description | Value |
| --- | --- | --- |
| $M$ | Number of rows | 5 |
| $N$ | Number of arcs | 4 |
| $\lambda$ | Radius of tuning curve | 1.6 |
| $U_{L4}/U_{L6}$ | Utilization rate of L4 / L6 synapses | 0.5 |
| $U_{ThC}/U_{L46}$ | Utilization rate of thalamocortical / L4-to-L6 synapses | 0.8 / 0.5 |
| $\alpha_{L4}/\alpha_{L6}$ | Slope of L4 / L6 gain function | 1 |
| $\theta_{L4}/\theta_{L6}$ | Threshold of L4 / L6 gain function | 5 / 3 |
| $J_{L4}^0/J_{L6}^0$ | Intra-column synaptic efficacy in L4 / L6 | 2.2 / 2.5 |
| $J_{L4}^1/J_{L6}^1$ | Vertical/horizontal inter-column synaptic efficacy in L4 / L6 | 0.05 / 0.03 |
| $J_{L4}^2/J_{L6}^2$ | Diagonal inter-column synaptic efficacy in L4 / L6 | 0.001 |
| $J_{ThC}$ | Synaptic efficacy of thalamocortical connection | 1 / 0.05 |
| $J_{L46}$ | Synaptic efficacy of L4-to-L6 connection | 0.24 |
| $\tau_m$ | Membrane time constant | 0.001 s |
| $\tau_{rec,L4}/\tau_{rec,L6}$ | Recovery time constant of L4 / L6 synaptic resources | 0.5 / 1 s |
| $\tau_{rec,ThC}/\tau_{rec,L46}$ | Recovery time constant of thalamocortical / L4-to-L6 synaptic resources | 0.8 / 1.2 s |

